## Supplemental Material for "Rapid dynamic naturalistic monitoring of bradykinesia in Parkinson’s disease using a wrist-worn accelerometer"

**Supplemental Material Naturalistic rapid PD bradykinesia monitoring Habets et al**

**Table S1**: Overview of extracted features. All features were extracted over x-, y-, z-, axes, and signal vector magnitude signals. Literature source shows first author name, see reference list for full reference.

| **Feature name** | **Source/ literature** |
|---|---|
| *Temporal domain* | |
| Maximum acc | Griffiths[1] |
| Interquartile range of acc | Griffiths[1] |
| 90th percentile of acc | Rispens [2] |
| median of acc | Hoff [3,] Keijsers [4] |
| mean of acc | Hoff [3,] Keijsers [4] |
| standard deviation of acc | Shawen [5] |
| coefficient of variation | Shawen [5] |
| variance of acc | Shawen [5] |
| acceleration range | Mahadevan [6] |
| low acc peaks (n) | Balasubramanian [7] |
| high acc peaks (n) | Balasubramanian [7] |
| time spent above 1g acc (%) | Keijsers [4,] Salarian [8] |
| acc entropy | Mahadevan [6,] Lonini [9] |
| jerkiness ratio/ smoothness | Mahadevan [6,] Lonini [9] |
| root mean square (RMS) | Mahadevan [6] |

| | |
|---|---|
| Ratio of x/y/z-RMS compared to vector magnitude-RMS | Sekine [10] |
| Axial cross-correlation (X-Y; (X-Z,; Y-Z) | Mahadevan [6,] Lonini [9] |
| *Spectral domain* | |
| spectral power < 3.5 Hz | Griffiths[1,] Evers [11] |
| spectral power 0.7 < 1.4 Hz | Griffiths[1,] Evers [11] |
| spectral power 1.4 < 2.8 Hz | Griffiths[1,] Evers [11] |
| spectral power 2.8 < 3.5 Hz | Griffiths[1,] Evers [11] |
| spectral flatness | Mahadevan [6] |
| spectral entropy | Mahadevan [6] |
| spectral variance | Balasubramanian [7,] Beck [12] |
| spectral smoothness | Balasubramanian [7] |
| spectral low/high peaks | Balasubramanian [7] |
| Dominant frequency magnitude | Mahadevan [6,] Lonini [9] |
| Dominant frequency ratio | Mahadevan [6] |
| Dominant frequency flatness | Mahadevan [6] |
| Dominant frequency entropy | Mahadevan [6] |

**Figure S1**: **Schematic visualization of data splitting method for individual models.**
Pre- and post-medication features were balanced in number.
During every fold in individual-model training and testing, ⅕ of pre-medication data and ⅕ of post-medication data (two times 10% of total data leads to 20% of total data), were selected as ‘test data’ for the validation of the model which was trained using the ‘training data’. For the training data selection, blocks of 2% of data adjacent to the test data were excluded, to decrease the temporal dependence of the training and test data.
Due to the expected difference in activities between blocks of 10% of data, we repeated model training and testing with every consecutive block of 10% of data. This led to 41 different training and testing folds. This figure visualizes folds 1, 2, 3, 18, 32, 40, and 41 as examples.

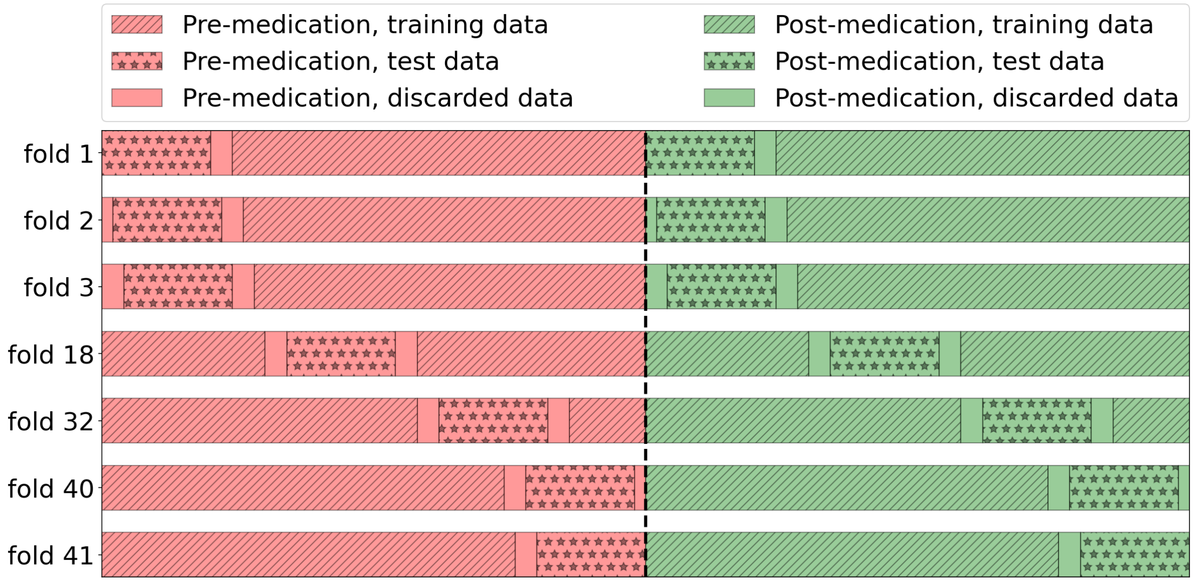

**Figure S2**: Visualization of activity filter performance versus the parallel raw signal vector magnitude.

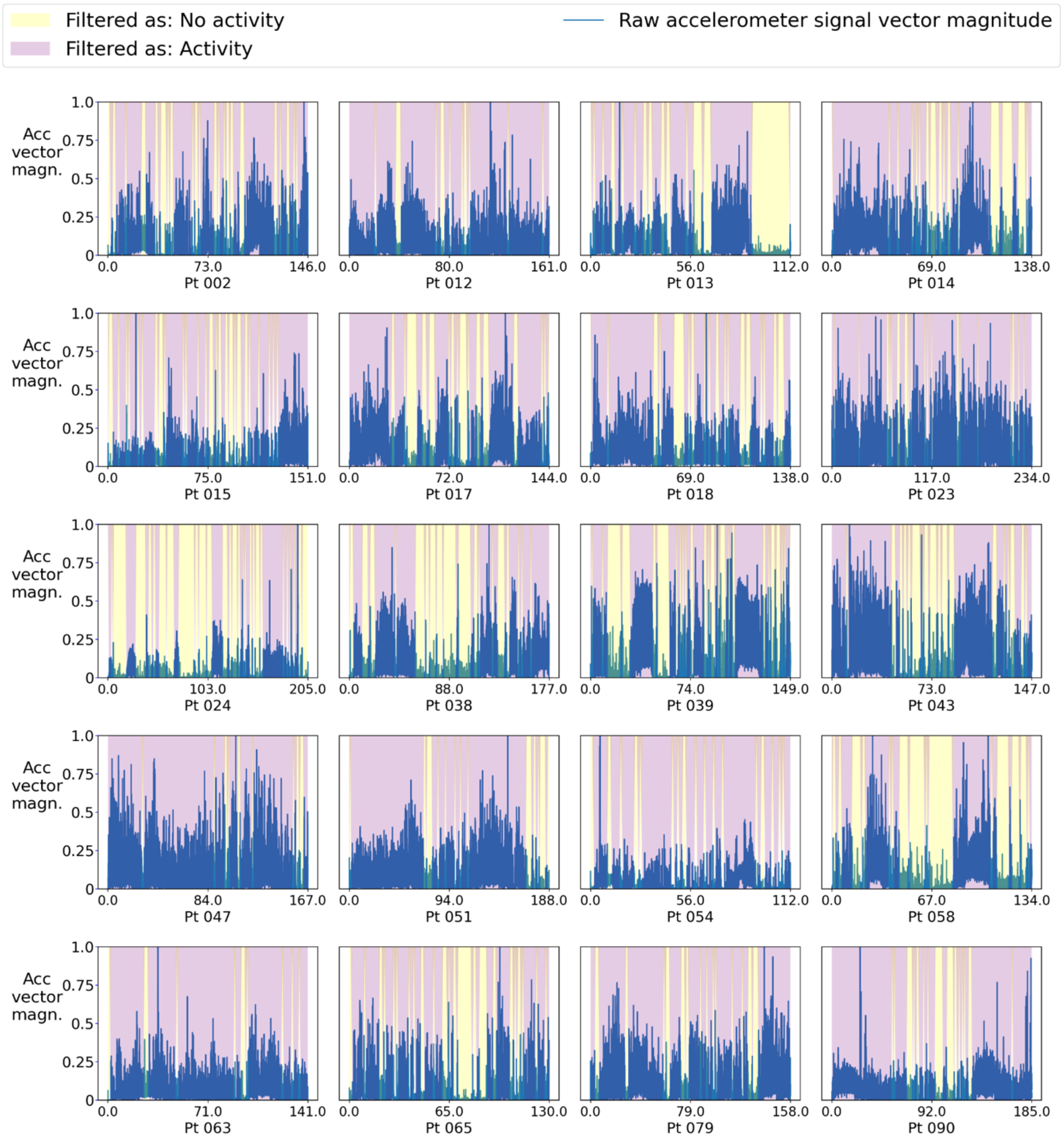

Y-axes represent the vector magnitude of the raw tri-axial accelerometer signal (in blue). The accelerometer vector magnitudes are individually normalized with a maximum of 1.0 for visualization purposes since maximum acceleration vectors vary between individuals.
X-axes represent time in minutes, where the first half consists of pre-medication recordings, and the second half of post-medication recordings.
The background color-shading represents whether the activity filtered classified the data epoch as activity present (purple), or not present (yellow).

**Table S2: Predictive metrics for all models and approaches**

| | | | All features | All features | 4 features | 4 features |
|---|---|---|---|---|---|---|
| *CLASSIFIER* | | n=20, mean (sd) | All minutes | Activity filtered | All minutes | Activity filtered |
| ***SUPPORT VECTOR*** | *INDIVIDUAL MODEL* | *auc* | 0.682 (0.15) | 0.696 (0.18) | 0.499 (0.12) | 0.533 (0.16) |
| | | *auroc, n sign* | 16 | 13 | 2 | 4 |
| | | *accuracy* | 0.632 (0.12) | 0.651 (0.14) | 0.490 (0.11) | 0.509 (0.14) |
| | | *accuracy, n sign* | 15 | 14 | 5 | 10 |
| | *GROUP MODEL* | *auroc* | 0.669 (0.10) | 0.703 (0.10) | 0.590 (0.11) | 0.633 (0.13) |
| | | *auroc, n sign* | 16 | 17 | 10 | 11 |
| | | accuracy | 0.624 (0.09) | 0.640 (0.08) | 0.560 (0.11) | 0.597 (0.08) |
| | | *accuracy, n sign* | 11 | 12 | 9 | 12 |
| ***RANDOM FOREST*** | *INDIVIDUAL MODEL* | *auroc* | 0.649 (0.13) | 0.656 (0.17) | 0.586 (0.12) | 0.619 (0.14) |
| | | *auroc, n sign* | 15 | 10 | 6 | 11 |
| | | *accuracy* | 0.611 (0.10) | 0.611 (0.13) | 0.558 (0.10) | 0.588 (0.11) |
| | | *accuracy, n sign* | 13 | 10 | 8 | 7 |
| | *GROUP MODEL* | *auroc* | 0.661 (0.10) | 0.698 (0.11) | 0.593 (0.11) | 0.636 (0.14) |
| | | *auroc, n sign* | 12 | 16 | 10 | 10 |

| | | | | | | | |
|---|---|---|---|---|---|---|---|
| | | *accuracy* | 0.598 (0.08) | 0.626 (0.08) | | 0.564 (0.08) | 0.587 (0.08) |
| | | *accuracy, n sign* | 12 | 12 | | 6 | 7 |

**Figure S3: Comparison of different model approaches for short window medication states classification**

To understand which methodological approaches yielded best performance, we explored the differences between individual and group trained models and the effect of an activity filter in detail and investigated optimal training data sizes and feature window lengths (see Methods section in main text). For this we compared the 20 areas under the receiver operator characteristic (AUC) and classification accuracy scores of each model using equality plots. Each dotted line visualizes the line x = y, and represents equality of the two displayed models.

The p-values throughout these figures indicate whether the ratio of patients that scored higher on model X versus model Y is statistically significant. We performed a 5000 permutation test where 20 dots (random x-value, random y-value) were randomly plotted in the equality plot. The p-values represent the chance that the distribution is better than the random chance level.

Figures S3A and S3B show classification superiority of models analyzing 60 seconds data epochs using 103 accelerometer-derived features compared to 4 features (AUC p-values < 0.002, accuracy p-values < 0.020, significance tested via a 5000-permutation test). The consecutive S3C-F figures only display the models including 103 features. Individual models based on a support vector classifier (SV) resulted in higher AUC scores and accuracies than individual models based on random forest classifier (RF) in 15 out of 20 patients (figure S3C, p = 0.009 below). SV and RF group models yielded similar AUC scores and accuracies (figure S3D below, p = 0.406). Overall, applying the activity filter led to slightly better mean results per model (table S2). On an individual level, there was no significant difference between classification performance with or without activity filtering (figure S3EF, p-values ranged between 0.06 and 0.41). However, it was noted that there was a trend towards higher individual predictive performance with activity filtering.

**Figure S3A**: Individual models: 4 versus 103 features.

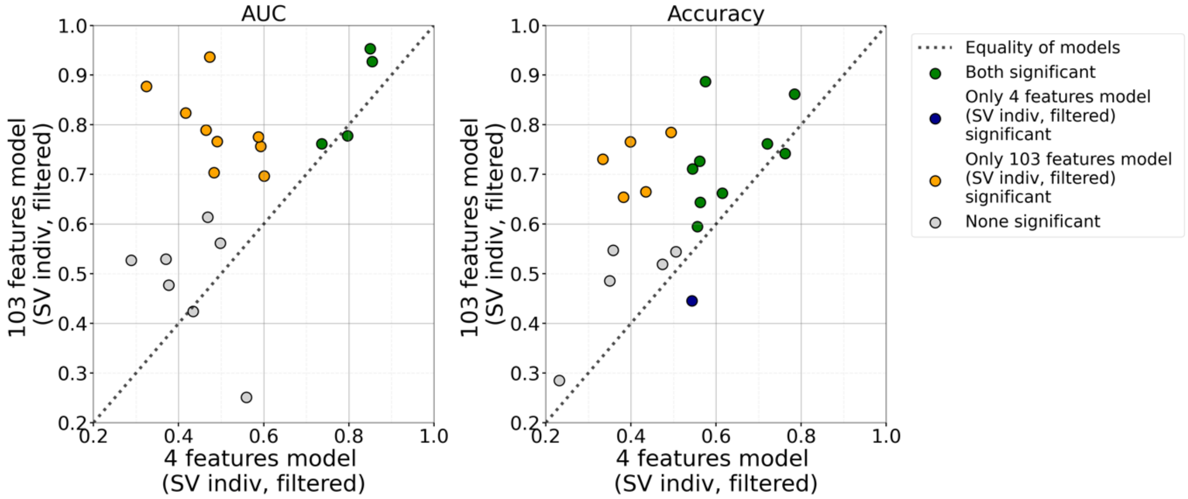

Individual 103 features model higher than individual 4 features model for:
AUC: 17 out of 20 higher, p < 0.000, accuracy: 18 out of 20 higher: p < 0.000

**Figure S3B**: Group models: 4 versus 103 features.

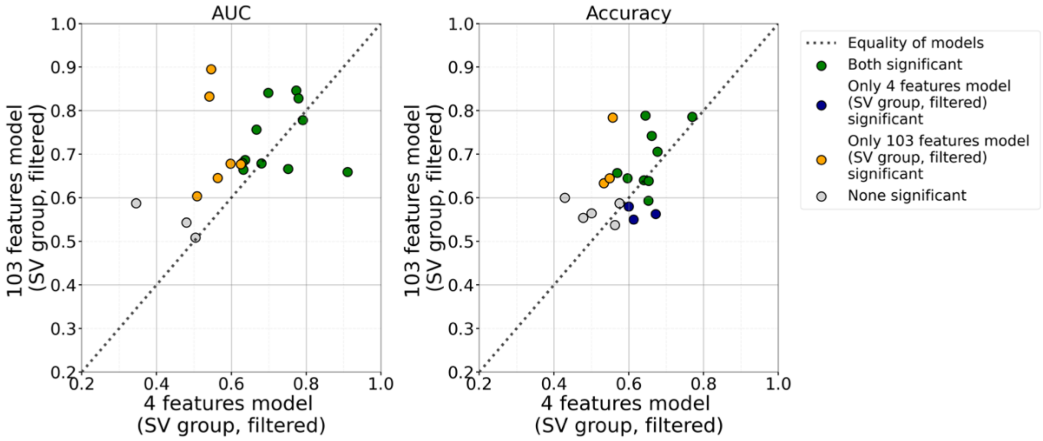

Group 103 features model higher than group 4 features model for:
AUC: 16 out of 20 higher, p = 0.002, accuracy: 14 out of 20 higher: p = 0.023

**Figure S3C**: Individual models: SV versus RF classifiers.

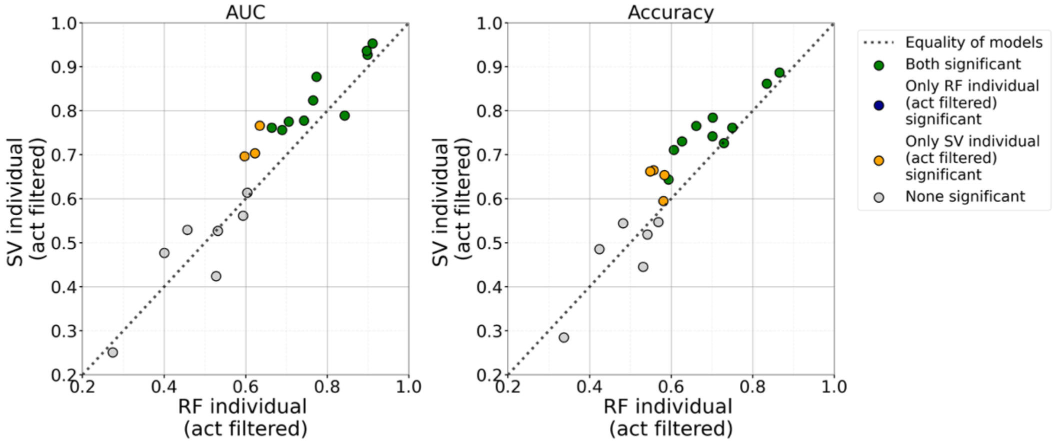

Individual SV model higher than individual RF model for:
AUC: 15 out of 20 higher, p = 0.009, accuracy: 15 out of 20 higher: p = 0.009

**Figure S3D**: Group models: SV versus RF classifiers.

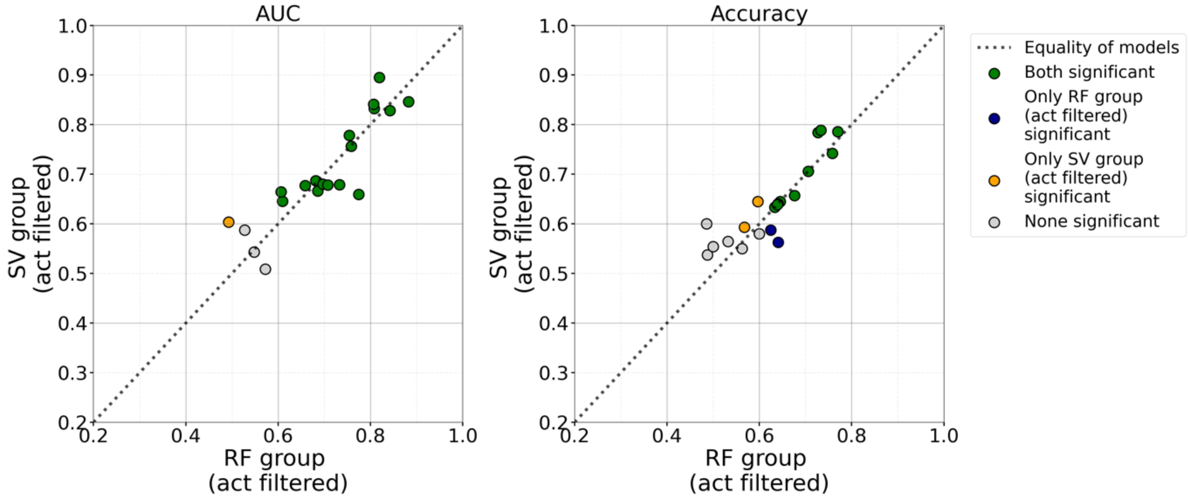

Group SV model higher than group RF model for:
AUC: 10 out of 20 higher, p = 0.406, accuracy: 10 out of 20 higher: p = 0.406

**Figure S3E**: Individual SV models: with activity filtering versus without activity filtering.

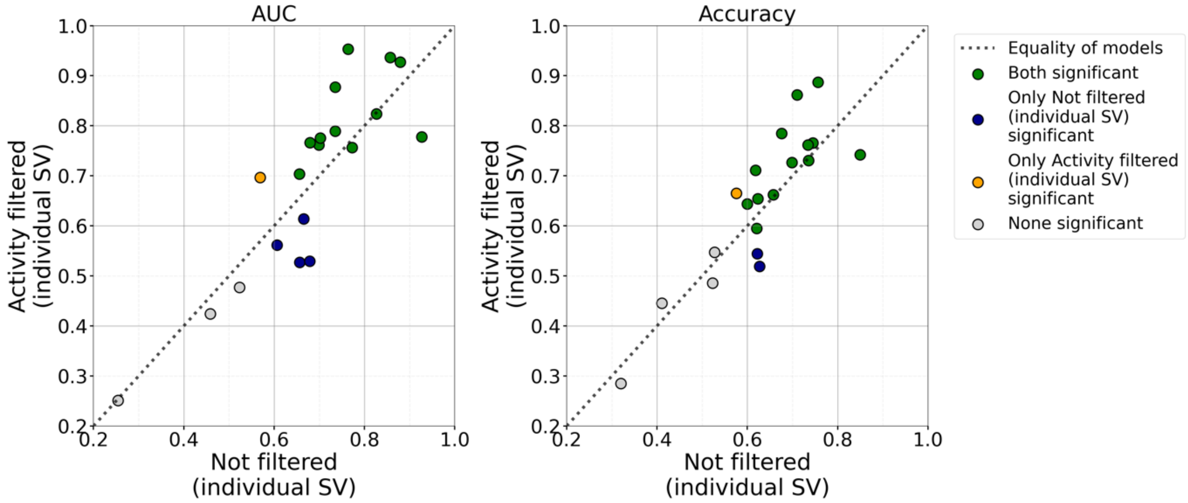

Individual SV model with activity filtering higher than individual SV model without activity filtering for:
AUC in 10 out of 20 higher, p = 0.406, accuracy in 13 out of 20 higher, p = 0.056.

Note that both the AUC scores and the classification accuracies of the activity filtered models are marginally higher than those of the not filtered models, when all ‘none significant’ candidates are disregarded. We conclude that although there is no statistically significant superiority of the activity filtered models, there is a trend that activity filtered models lead to higher predictive performance and this step was included within our standard pipeline.

**Figure S3F**: Group SV models: with activity filtering versus without activity filtering.

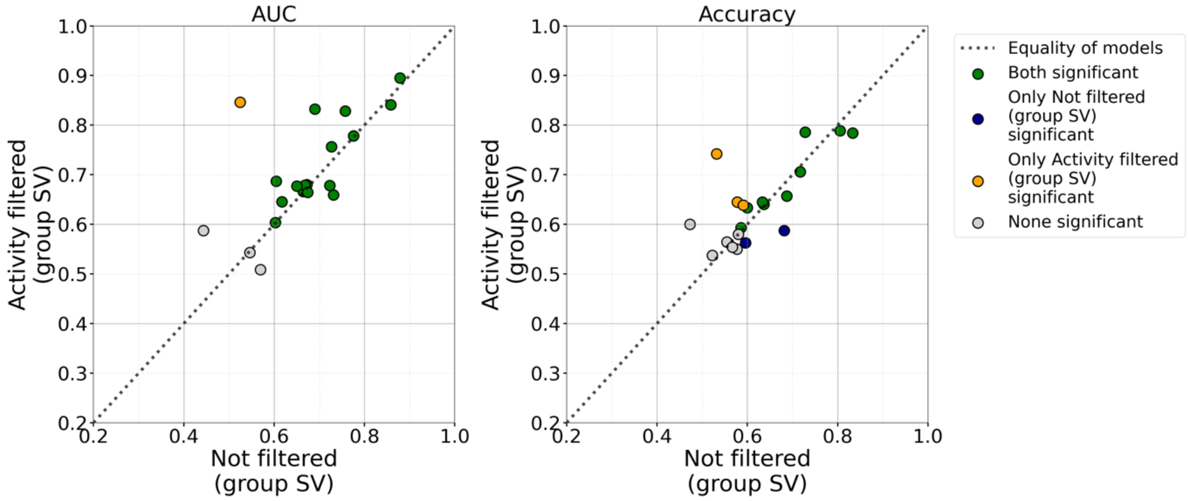

Group SV model with activity filtering higher than group SV model without activity filtering for:
AUC in 13 out of 20 higher, p = 0.056, accuracy in 11 out of 20 higher, p = 0.250.

**Figure S4A: Good classification performance in patients with and without tremor**

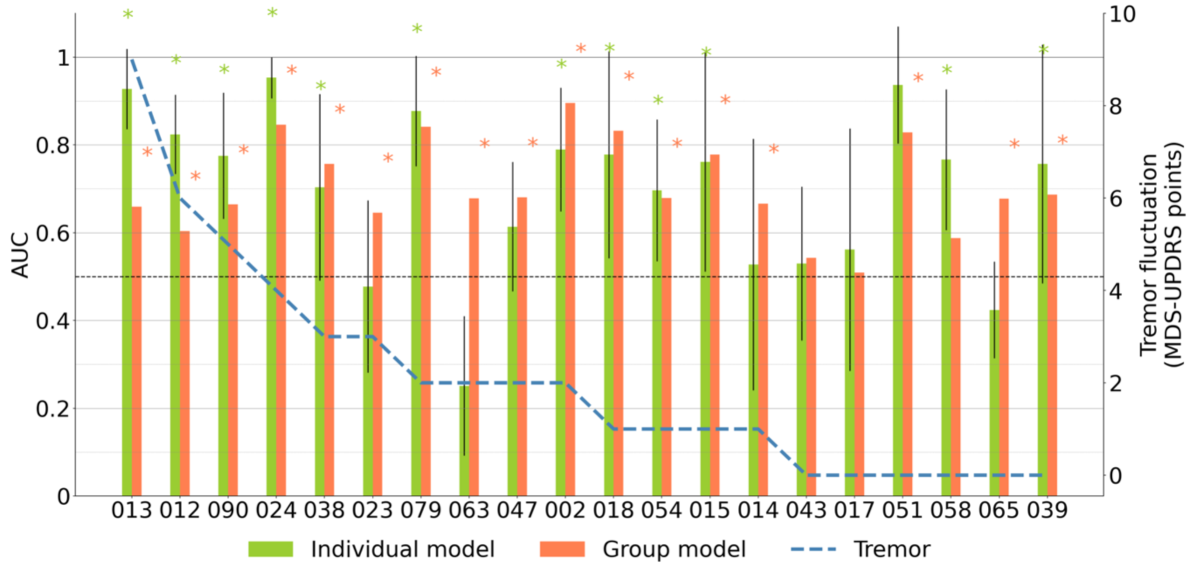

**Figure S4B: Good classification performance in patients with and without abnormal involuntary movements**

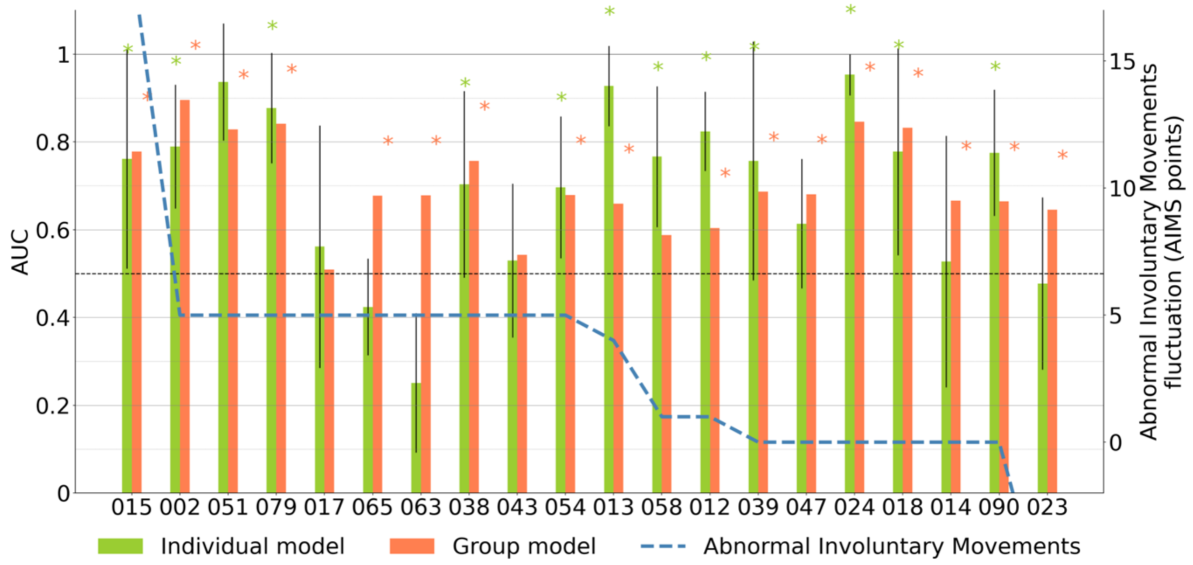

Colored bars visualize the individual AUC scores from the best individual and the best group model (both support vector classifier, and activity filtered), and correspond to the left y-axis. Individual tremor fluctuations between pre- and post-medication correspond to the right y-axis. Tremor scores represent the described MDS-UPDRS III items for unilateral upper extremity tremor (see Methods). On the x-axis individual participants are sorted on tremor fluctuation, in descending order. Colored asterisks indicate statistical significance of the AUC score compared to chance level (alpha = 0.05, FDR corrected). The black dotted line indicates chance-level for the AUC scores. AUC: area under the receiver operator characteristic; FDR: false discovery rate, MDS-UPDRS: Movement Disorders Society - Unified Parkinson Disease Rating Scale.

**Table S3: Spearman R correlations between symptom fluctuation and predictive performance at an individual level.** Spearman r correlations are calculated between the MDS-UPDRS tremor and bradykinesia and AIMS fluctuations (collected and calculated as described in the Methods), and the predictive performance per participant. Support vector models including activity filtering were compared for individual and group model comparisons.

| | Individual models, AUC | Group models, AUC | Individual models, accuracy | Group models, accuracy |
|---|---|---|---|---|
| Bradykinesia (r (p)) | 0.24 (p = 0.305) | 0.01 (p = 0.962) | 0.24 (p = 0.305) | 0.18 (p = 0.452) |
| Tremor (r (p)) | 0.34 (p = 0.140) | 0.11 (p = 0.642) | 0.21 (p = 0.380) | -0.06 (p = .807) |
| Abnormal involuntary movements (r (p)) | -0.05 (p = 0.851) | 0.122 (p = 0.607) | 0.01 (p = 0.979) | 0.34 (p = 0.142) |
